## Extended figures for "AlphaPeptDeep: A modular deep learning framework to predict peptide properties for proteomics"

**Extended Figure 1. Amino acid (AA) and modification embedding.**

There are two ways to embed AAs in AlphaPeptDeep: (1) one-hot encoding and (2) *torch.Embedding*. As shown in the left panel, one-hot encoding in AlphaPeptDeep generates a one-hot vector that is filled with zeros except for the position indicated by ID of the AA, which is set to one. The one-hot vector is then mapped into an embedded vector (also called tensor) using a linear layer. *torch.Embedding* generates an embedded vector directly from the ID of each AA.

To represent a modification on an AA site, we use its chemical composition. To generate the embedded vector, we fix the number of C, H, N, O, S, and P atoms as the first 6 elements of the vector. For other atoms, we map them into a 2-D vector using a linear layer and concatenate the 2-D vector with the fixed 6 elements. Finally, a vector of size 8 is able to represent a modification.

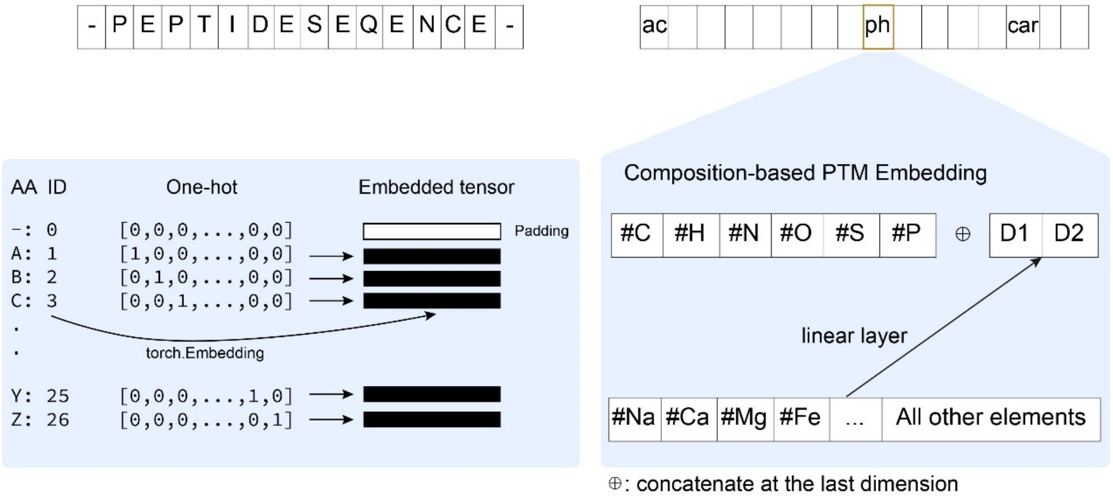

**Extended Figure 2. An illustrative example of using the model shop module.**

We built a deep learning model with only a few lines of code to predict if a peptide elutes in the first half or the second half of the LC gradient ('is_first_half_lc'). We then called 'model.train()' on the training set to train the model and 'model.predict()' on the testing set. We used the 351,804 PSMs of tryptic HeLa peptides identified by MaxQuant at 1% FDR measured by a two-hour gradient[1]. 'is_first_half_lc' is set as 1 if the PSM is at the first hour, otherwise 0. We then split the PSMs into two sets, where 80% for training and 20% for testing, without any intersections on sequence-level. The training took 16 min, and achieved 95% accuracy on the 70,018 testing PSMs.

```
from peptdeep.model.model_shop import (
    Model_for_Generic_ModAASeq_BinaryClassification_Transformer,
    ModelInterface_for_Generic_ModAASeq_BinaryClassification
)

model = ModelInterface_for_Generic_ModAASeq_BinaryClassification(
    model_class=Model_for_Generic_ModAASeq_BinaryClassification_Transformer
)
model.target_column_to_train = 'is_first_half_lc'
model.target_column_to_predict = 'predicted_is_first_half_lc'
model.train(train_df, epoch=50, warmup_epoch=20, lr=1e-5, verbose=True)
model.predict(test_df)
```

✓ 16m 13.6s

| | sequence | mods | mod_sites | rt_norm | is_first_half_lc | predicted_is_first_half_lc |
|---|---|---|---|---|---|---|
| 0 | DFSHTGR | | | 0.103493 | 1 | 0.998056 |
| 8 | RLPPIVR | | | 0.343450 | 1 | 0.940949 |
| 13 | DYHATWK | | | 0.230220 | 1 | 0.997083 |
| 14 | ISHFTQR | | | 0.127492 | 1 | 0.997336 |
| 15 | ISHFTQR | | | 0.127609 | 1 | 0.997336 |
| ... | ... | ... | ... | ... | ... | ... |
| 351797 | SPVGSGAPQAAAPAPAAHVAGNPGGDAAPAATGTAAAASLATAAGS... | | | 0.644790 | 0 | 0.005870 |
| 351798 | AAAAAAAAAAAATGTEAGPGTAGGSENGSEVAAQPAGLSGPAEVGP... | | | 0.718966 | 0 | 0.001388 |
| 351799 | SPVGSGAPQAAAPAPAAHVAGNPGGDAAPAATGTAAAASLATAAGS... | | | 0.624617 | 0 | 0.005870 |
| 351800 | AAAAAAAAAAAATGTEAGPGTAGGSENGSEVAAQPAGLSGPAEVGP... | | | 0.718805 | 0 | 0.001388 |
| 351801 | SPVGSGAPQAAAPAPAAHVAGNPGGDAAPAATGTAAAASLATAAGS... | | | 0.630824 | 0 | 0.005870 |

70018 rows × 6 columns

Testing accuracy = 95% (pred cutoff=0.5)

**Extended Figure 3. Prediction speed of the MS2, RT and CCS models in AlphaPeptDeep.**

Peptides were in silico digested from reviewed E. coli, fission yeast, and human protein sequences by using trypsin with at most 2 missed cleavages, with peptide lengths from 7 to 30. Oxidation@M was set as a variable modification with a maximum occurrence of 2. (a) Overall peptide and precursor (peptide with charge) numbers for each species. (b) prediction time on CPU with multiprocessing. There is no separate time for CCS prediction as CCS and MS2 are predicted together in the multiprocessing mode. (c) prediction time on a RTX3080 GPU. Note that even for predicting nearly 8M precursors from human proteome, the total prediction time for RT, CCS, and MS2 was less than 10 minutes. Saving the predicted values into the HDF file took only ~2 minutes. This demonstrates that generating a predicted spectral library is very fast if a GPU is used. When multiprocessing is used, the prediction time is acceptable (~2 hours).

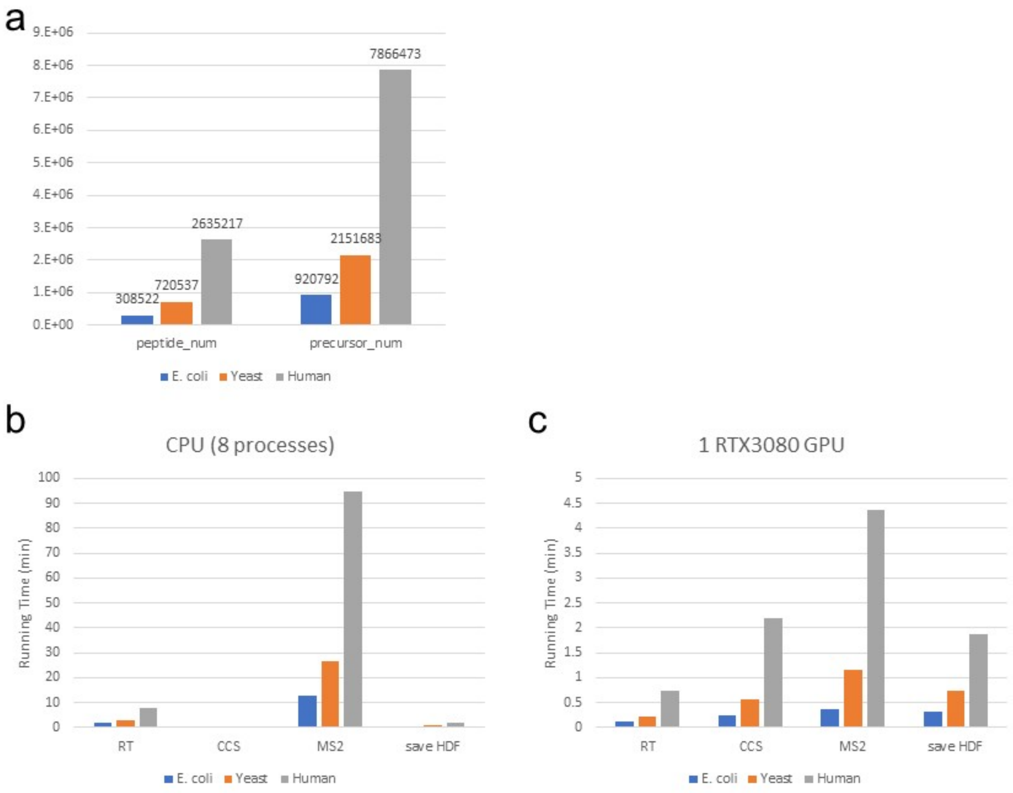

**Extended Figure 4. Comparing transformers with LSTM.**

The transformer and LSTM MS2 prediction models were trained and tested on the same tryptic peptides. The LSTM model is very similar to the pDeep model that we used before. The comparison shows that the transformer is slightly better than LSTM for MS2 prediction.

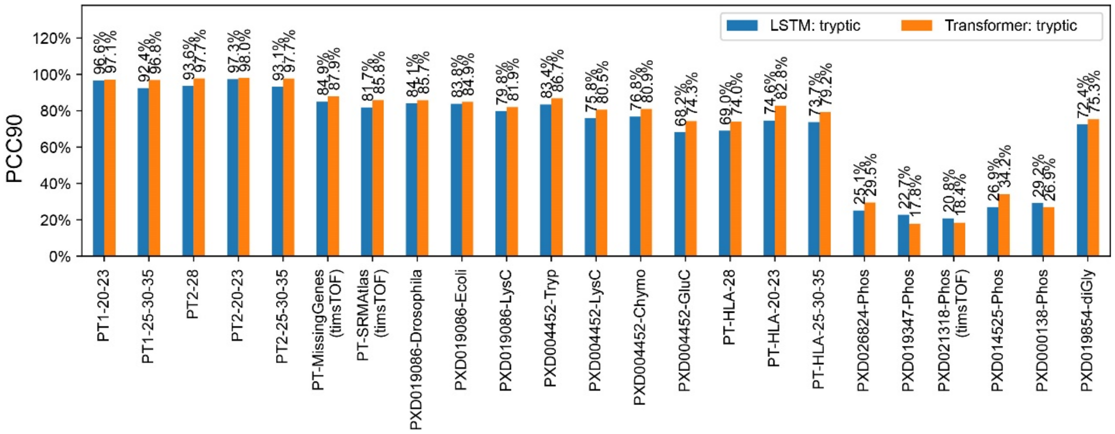

**Extended Figure 5. RT model fine-tuning for short LC gradients.**

Fine-tuning the RT model on **(a)** 30- and **(b)** 15-minute LC gradient. MaxQuant results were downloaded from PXD006932.[2]

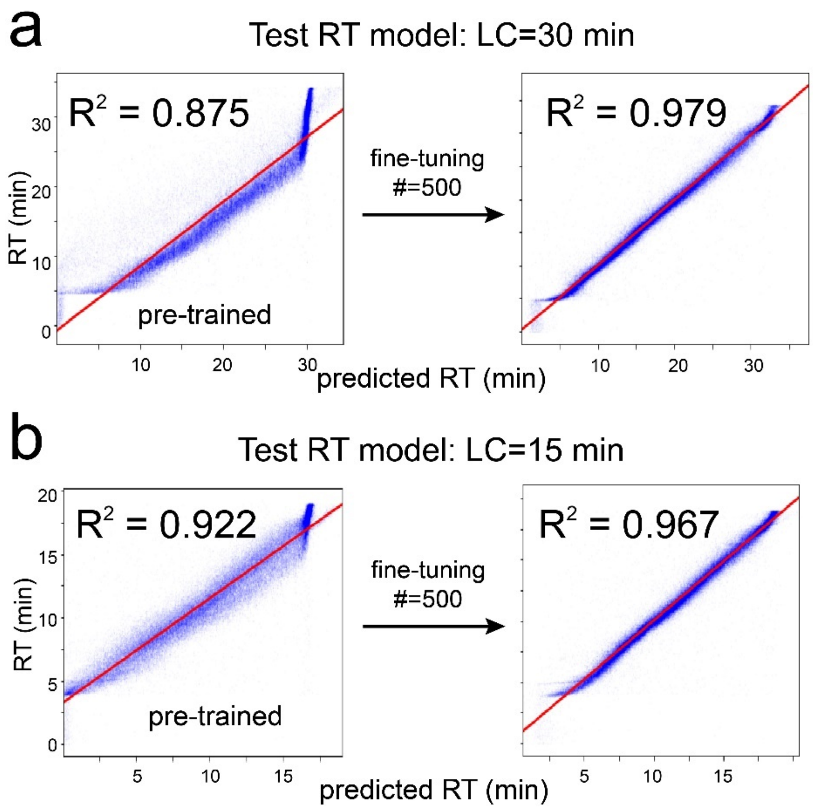

**Extended Figure 6. Visualizing transformer model attention with bertviz.**

The attention values of the 8 heads in the first transformer layer (Layer 0) and the last layer (Layer 3) of the pre-trained and transfer-learned MS2 models are shown. The right panels are the zoom-in plots of head 2 and 3 of these two models.

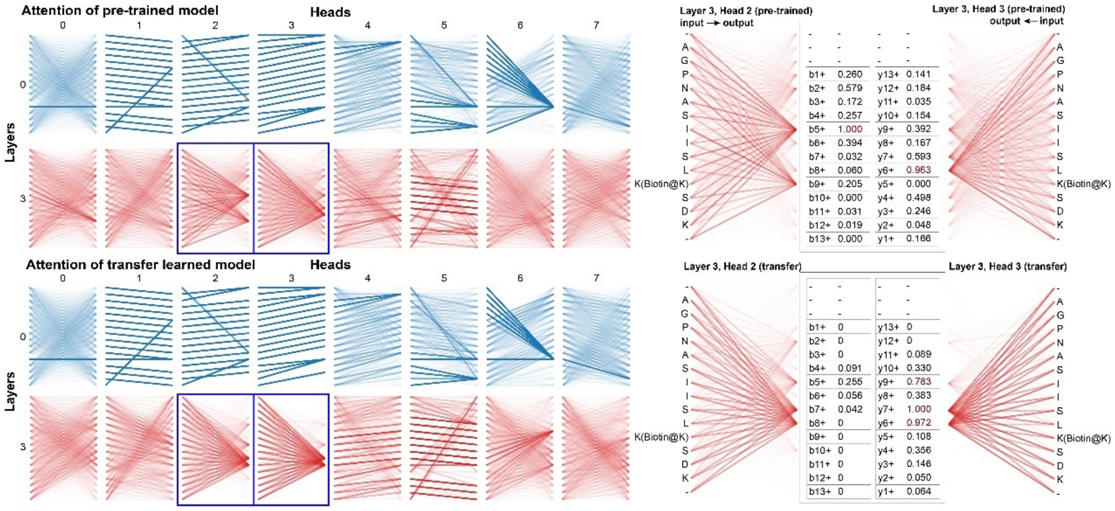

**Extended Figure 7. Sequence motifs of identified phospho-HLA peptides**.

Unmodified and phosphorylated peptides with 9-, 10-, and 11-mers are displayed. The peptides in the first panel were identified on tumor dataset from one of the patients (Mel-15). Peptides from other panels were identified from mono-allelic dataset.

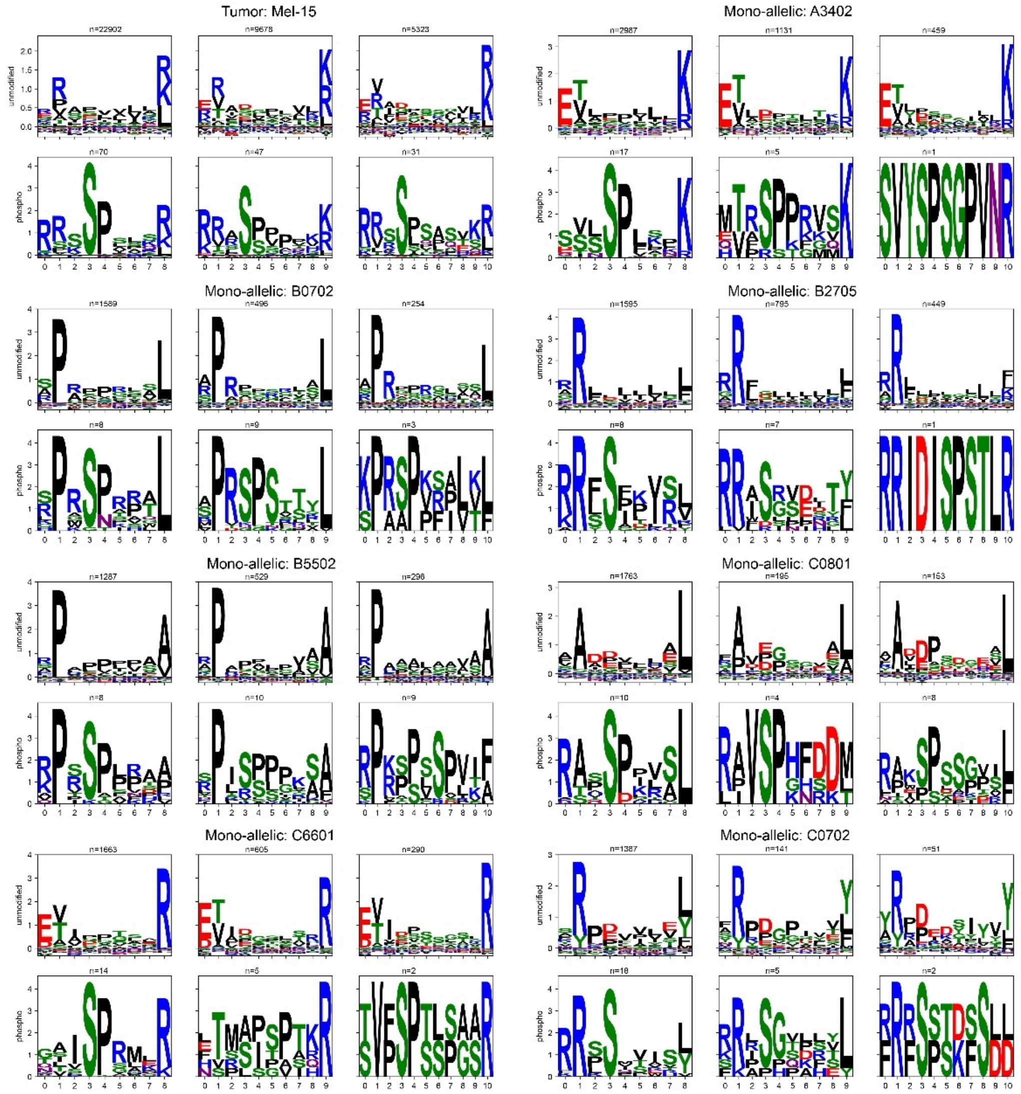

**Extended Figure 8. Inspecting of phospho-HLA peptides using MS2 and RT prediction.**

PCC distribution between experimental and predicted MS2 spectra for unmodified and phospho-HLA peptides on **(a)** the mono-allelic and **(b)** tumor datasets. It shows that PCCs of phospho-HLA peptides are quite close to the unmodified ones. The RT difference between detected and predicted RT values are displayed in **(c)** and **(d)** on the mono-allelic and tumor datasets, respectively.

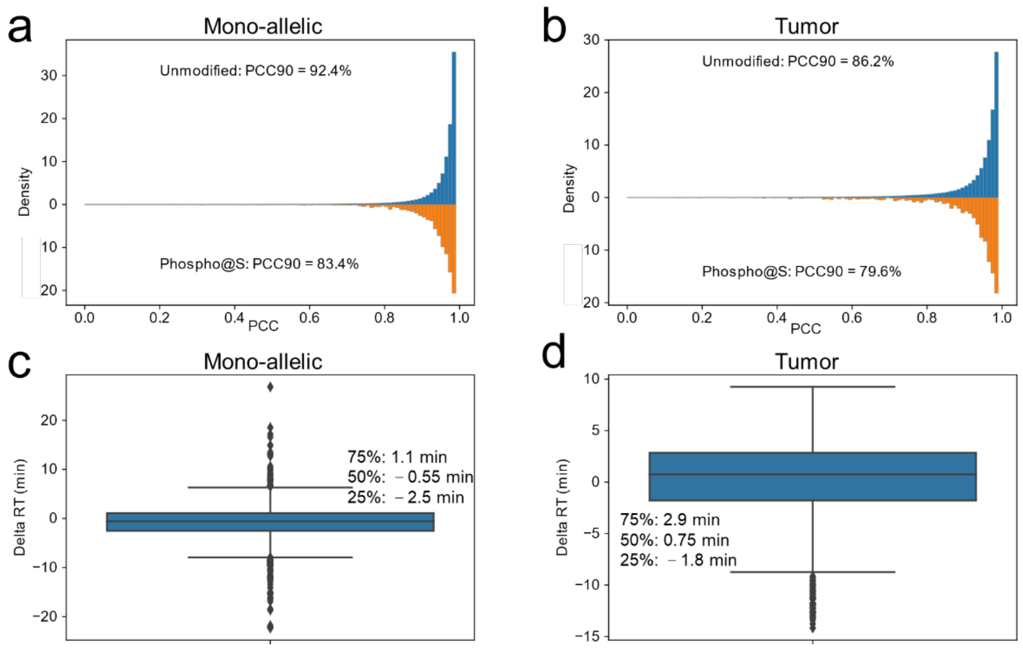

**Extended Figure 9. Identified modifications by AlphaPeptDeep with Open-pFind.**

(a) Mono-allelic dataset. (b) Tumor dataset.

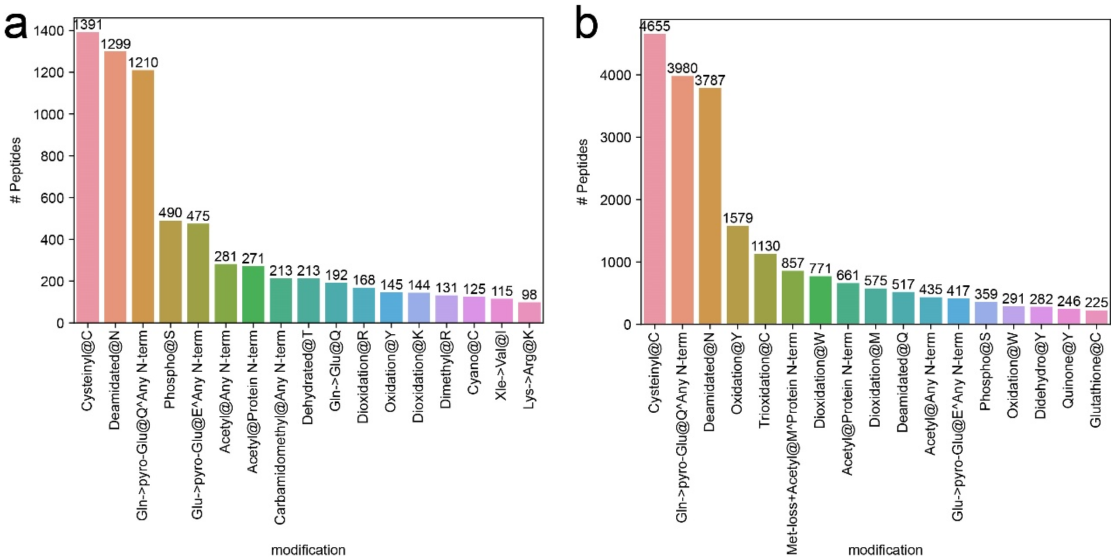

**Extended Figure 10. Producer-consumer schema for rescoring with multiprocessing**

The rectangle in the middle represents the main process, which contains model fine-tuning, prediction step on GPU, and the final Percolator rescoring. Other steps are processed in parallel, such as fragment matching, MS2 similarity calculation, and other feature extraction for Percolator rescoring. SA refers to spectral angle, and SPC refers to Spearman's rank correlation.

AlphaPeptDeep re-scores PSMs using a specified process number (8 by default).

All raw files are scheduled by the producer-consumer schema. After deep learning models are fine-tuned, the main process puts all RAW files in a queue and pops the first one into the producer processes for fragment extraction once there is an idle producer. The data will then be sent back to the main process for prediction, and this step is very fast using GPU. Next, the matched and predicted fragments will be sent to the consumer processes to extract all the 61-machine learning (ML) features. After all RAW files are processed, the PSMs and their ML features are combined in the main process for Percolator rescoring. From the perspective of each RAW file, it appears to be processed continuously without waiting for other RAW files.

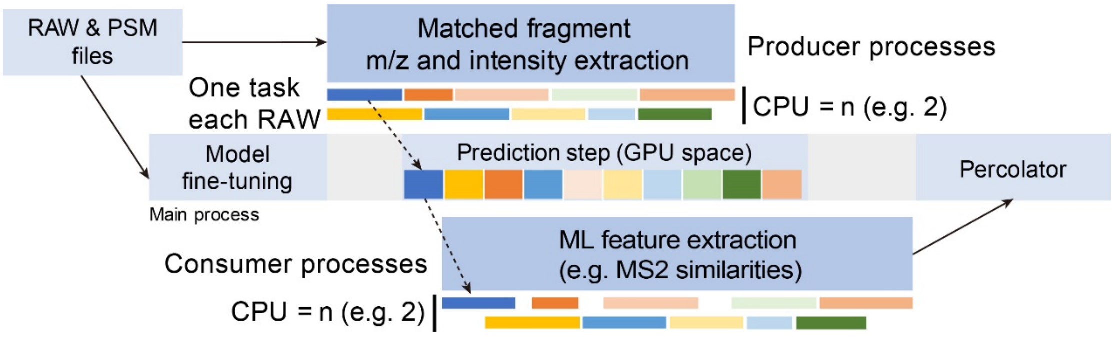
